# Individual differences in volitional social motivation in male and female mice following social stress

**DOI:** 10.1101/2022.11.08.515718

**Authors:** Jovana Navarrete, Kevin N. Schneider, Briana M. Smith, Nastacia L. Goodwin, Yizhe Y. Zhang, Ethan Gross, Valerie S. Tsai, Mitra Heshmati, Sam A. Golden

**Author notes:** Correspondence to: Sam A. Golden. Denotes equivalent contribution (Jovana Navarrete, Kevin N. Schneider).

## Abstract

**Background:** A key challenge in developing new treatments for neuropsychiatric illness is the disconnect between preclinical models and the complexity of human social behavior. We aimed to integrate voluntary social self-administration into a preclinical rodent stress model, as a platform for the identification of basic brain and behavior mechanisms underlying stress-induced individual differences in social motivation. Here, we introduce an operant social stress (OSS) procedure with male and female mice, where lever presses are reinforced by freely moving social interaction with a familiar social partner across social stress exposure.

**Methods:** OSS is composed of three phases: (***i***) social self-administration training, (***ii***) social stress concurrent with daily reinforced social self-administration testing, and (***iii***) post-stress operant social reward testing under both non-reinforced and reinforced conditions. We resolve social stress-induced changes to social motivation behaviors using hierarchical clustering and aggregated z-scores, capturing the spectrum of individual differences that we describe with a social index score.

**Results:** OSS captures a range of stress-related dynamic social motivation behaviors inclusive of sex as a biological variable. Both male and female mice lever press for access to a social partner, independent of social partner coat color or familiarity. Social stress attenuates social self-administration in males and promotes social reward seeking behavior in females. Hierarchical clustering does not adequately describe the relative distributions of social motivation following stress, which we find is better described as a non-binary behavioral distribution that we define by introducing the social index score. This index is stable across individual mice.

**Conclusion:** We demonstrate that OSS can be used to detect stable individual differences in stress-induced changes to social motivation in male and female mice. These differences may reflect unique neurobiological, cellular and circuit mechanisms not captured by preclinical models that omit voluntary social behaviors. The inclusion of volitional social procedures may enhance the understanding of behavioral adaptations promoting stress resiliency and their mechanisms under more naturalistic conditions.

## Introduction

An ongoing focus of behavioral neuroscience is the identification of mechanisms that govern individual responses to social stress. Decades of work indicate that diverse neuronal and non-neuronal mechanisms drive this variability^1–3^. Even though coping strategies vary widely between individuals and play a significant role in subsequent vulnerability to neuropsychiatric disease^4, 5^, little is known about how coping strategies for social stress integrate features like social motivation. Further, even though many people (greater than 50% of the general population) experience significant social and emotional stress in their lives, only a subset of individuals (less than 10%) develop stress-related illness^6, 7^. Of these, women are more likely than men to be diagnosed with major depressive disorder and anxiety disorders, and the selection of coping strategy differs between sexes^8–11^. Since the selection of coping strategies vary widely, expanding preclinical stress models to include volitional social motivation may better capture the underlying individual neurobiological differences engaged by social stress.

A critical component of responding to stress is the decision to actively engage, or avoid, socially rewarding interactions. There is strong evidence that mice will voluntarily lever press or nose-poke for access to social partners, both for the reinforcing effects of aggressive^12–17^ and affiliative^18–20^ social interaction. The addition of operant social self-administration procedures to preclinical neuropsychiatric rodent models has revealed unexpected results, like the existence of compulsive addiction-like aggression seeking^17^ and the role of voluntary social interaction in blunting incubation of drug craving^21^. However, such voluntary social self-administration procedures have not yet been incorporated into social stress models, which predominantly rely on involuntary social interaction tests. Here, we introduce operant social stress (OSS), a behavioral procedure to evaluate voluntary social motivation in both male and female mice before, during and after repeated social stress exposure. OSS is composed of three phases: (***i***) social self-administration training, (***ii***) social stress concurrent with daily reinforced self-administration to test social decision-making, and (***iii***) operant social reward seeking under both non-reinforced and reinforced conditions (Figure 1).

**Figure 1.**
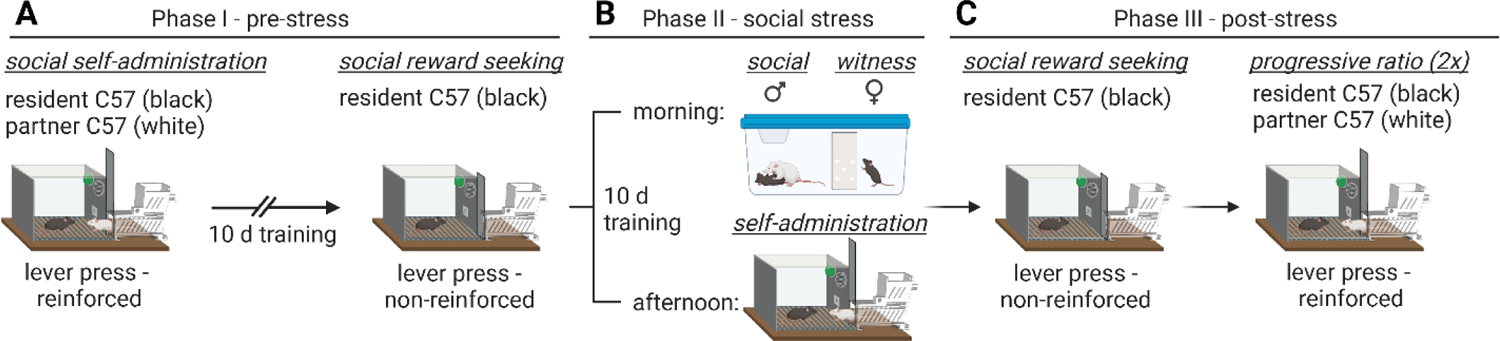
Operant social stress procedure schematic. (**A**) Phase I (pre-stress): male and female mice were trained for 10-d to lever press for freely moving access to a familiar social partner, followed by a 1-hr non-reinforced reward seeking test. (**B**) Phase II (social stress exposure): mice that acquired social self-administration were exposed to either social defeat (males) or witness defeat (females) for 10 d, paired with daily self-administration test sessions with a familiar social partner. (**C**) Phase III (post-stress): following the final social stress exposure day, mice were tested for social reward seeking a under extinction conditions (left), followed by 2 days of progressive ratio testing (right) to measure their social motivation breakpoint.

Currently, preclinical investigations of social stress behavior primarily use procedural variations of social defeat or witness defeat stress. In these procedures, mice are either subjected to repeated antagonistic social encounters^22–26^ or witness these repeated encounters through sensory but not physical contact^27, 28^. Following repeated defeats, mice are then evaluated in an involuntary social interaction test that measures their exploratory sensory contact with a non-familiar target mouse^26^. While clinical susceptibility and resilience exist on a spectrum^29–32^, preclinical defeat procedures use an arbitrary binary classification for stress phenotypes based on this sensory contact duration. Although social and witness defeat procedures offer the significant benefits of high-throughput and standardization, the inclusion of social self-administration procedures concurrent with social stress exposure will (*i*) allow assessment of voluntary social motivation, (*ii*) provide expanded measures of individual variability in voluntary social behavior, and (*iii*) provide a behavioral platform for resolving their underlying mechanisms.

In a series of experiments, we combine previously established models of social stress^26, 27, 33^ with mouse social self-administration procedures^17–20^ and operant reward testing to investigate volitional social stress responses. The results of our first set of experiments demonstrate that inbred male and female mice reliably acquire social self-administration of a partner, regardless of coat color or prior social familiarity, and exhibit robust social seeking behavior under non-reinforced conditions. Male mice exposed to social defeat stress exhibit attenuated social self-administration and social motivation compared to non-stressed controls, while female mice exposed to witness defeat stress exhibit enhanced social motivation compared to non-stressed controls. We then use these operant data to create a social index that describes individual differences in social motivation following stress. In this way, the OSS procedure provides a flexible behavioral platform for identifying fundamental mechanisms that drive individual responses to stress-induced changes in voluntary social motivation.

## Main Manuscript Materials and Methods

A detailed description of experimental subjects, apparatus and procedures are provided in the Supplementary Methods. Below, we briefly describe the five specific experiments:

### The operant social stress (OSS) procedure

Operant social stress (OSS) is composed of three phases: (*i*) social self-administration training and reward seeking, (*ii*) social stress exposure concurrent with social self-administration testing, and (*iii*) post-stress operant social reward testing under both non-reinforced and reinforced conditions (Figure 1). In Phase I, experimental male and female black-coated C57BL/6J mice are trained to press a lever for freely moving physical access to a familiar same-sex conspecific partner (white-coated C57BL/6J for compatibility with behavioral annotation) in a trial-based, 12-trial/day design for 10 days (Figure 1A and 2A). Following social self-administration acquisition, social reward seeking is measured using a 1-hr non-reinforced reward seeking test under extinction conditions. In Phase II, male mice are exposed to social defeat stress and female mice to witness defeat stress, once daily for 10 days, and 4 hr after each daily stress exposure they undergo social self-administration testing with their social partner (Figure 1B and 3A, left). Lastly, in Phase III, experimental mice are tested for non-reinforced social reward seeking and reinforced social progressive ratio responding (Figure 1C and 3A, right). These operant metrics are used to stratify mice according to social stress-induced dysregulation of social self-administration and reward seeking (Figures 4-6).

**Figure 2.**
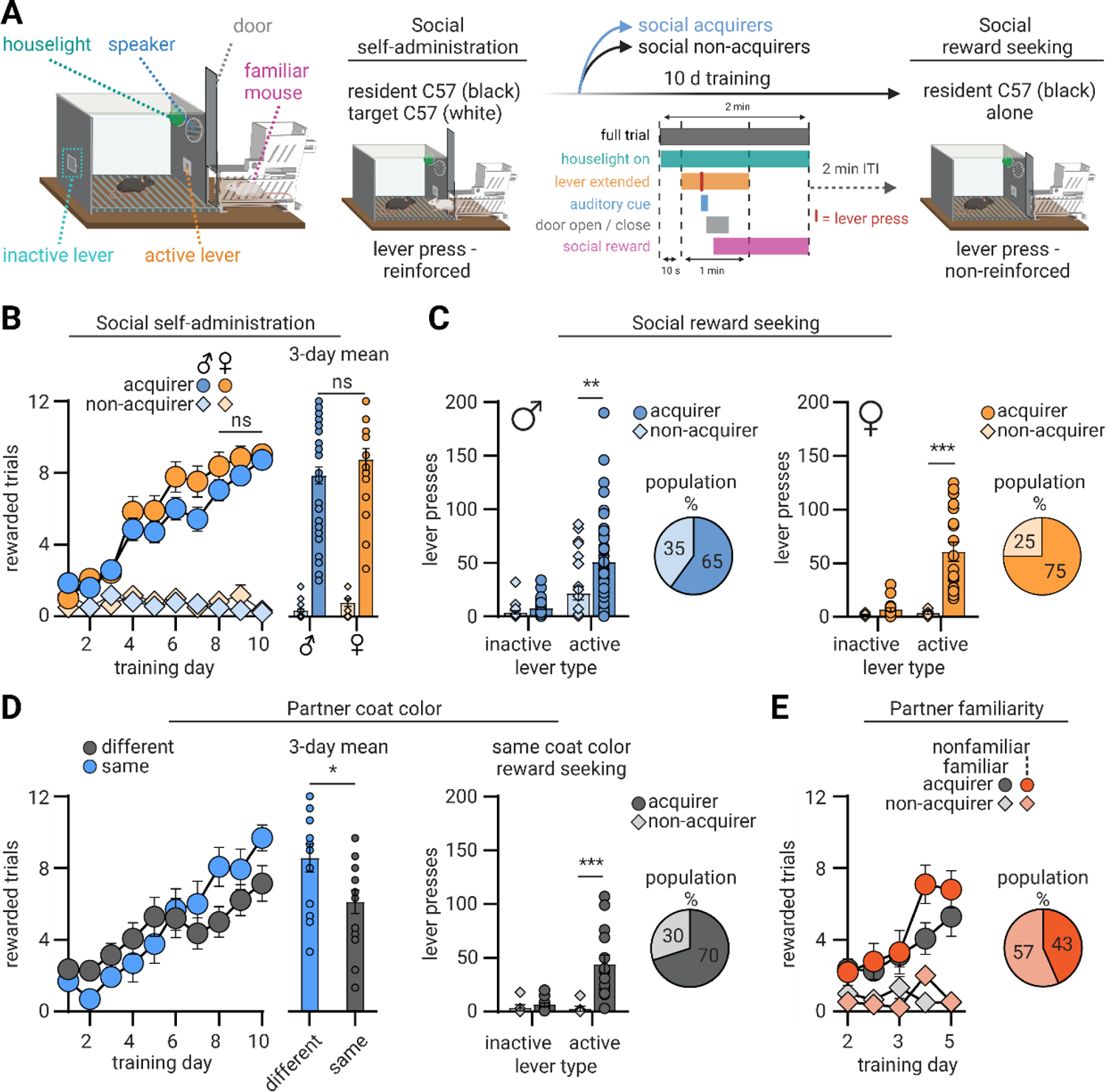
Male and female mice similarly acquire social self-administration an exhibit social reward seeking. (**A**) OSS chamber (left) and experimental timeline schematic (right) for social self-administration and social reward seeking. Resident male and female C57Bl/6J mice were trained to lever-press for access to a familiar social partner. In the individual trial schematic, the vertical red line within the ‘lever extended’ bin indicates an active lever press and the resulting sequence of events. (**B**) <u>Left</u>: Number of rewarded trials over 10 days (48-minute session/day; 12 trials/day) of social self-administration under a trial-based fixed-ratio 1 reinforcement schedule with male (n=67) and female (n=24). <u>Right</u>: Mean rewarded trials from the final 3-d of social self-administration. (**C**) Number of non-reinforced male (left) and female (right) active and inactive lever presses during a 1-hr social reward seeking test under extinction conditions following social self-administration. Pie charts show the proportion of each cohort that acquired social self-administration. (**D**)<u>Left</u>: Number of rewarded trials over 10 days (12 trials/day) of social self-administration with similar (n=20) and different (n=24) coat color social partners. <u>Middle</u>: Mean rewarded trials from the final 3-d of social self-administration. <u>Right</u>: Number of non-reinforced active and inactive lever presses during a 1-hr social reward seeking test under extinction conditions following social self-administration with a same coat color social partner, pie chart shows percent of mice that acquired social self-administration. (**E**) Number of rewarded trials over 10 days (12 trials/day) of social self-administration with familiar (n=20) and non-familiar (n=23) social partners, pie chart shows percent of mice that acquired social self-administration. Individual data denoted with symbols. *p < .05. Data are mean ± SEM.

**Figure 3.**
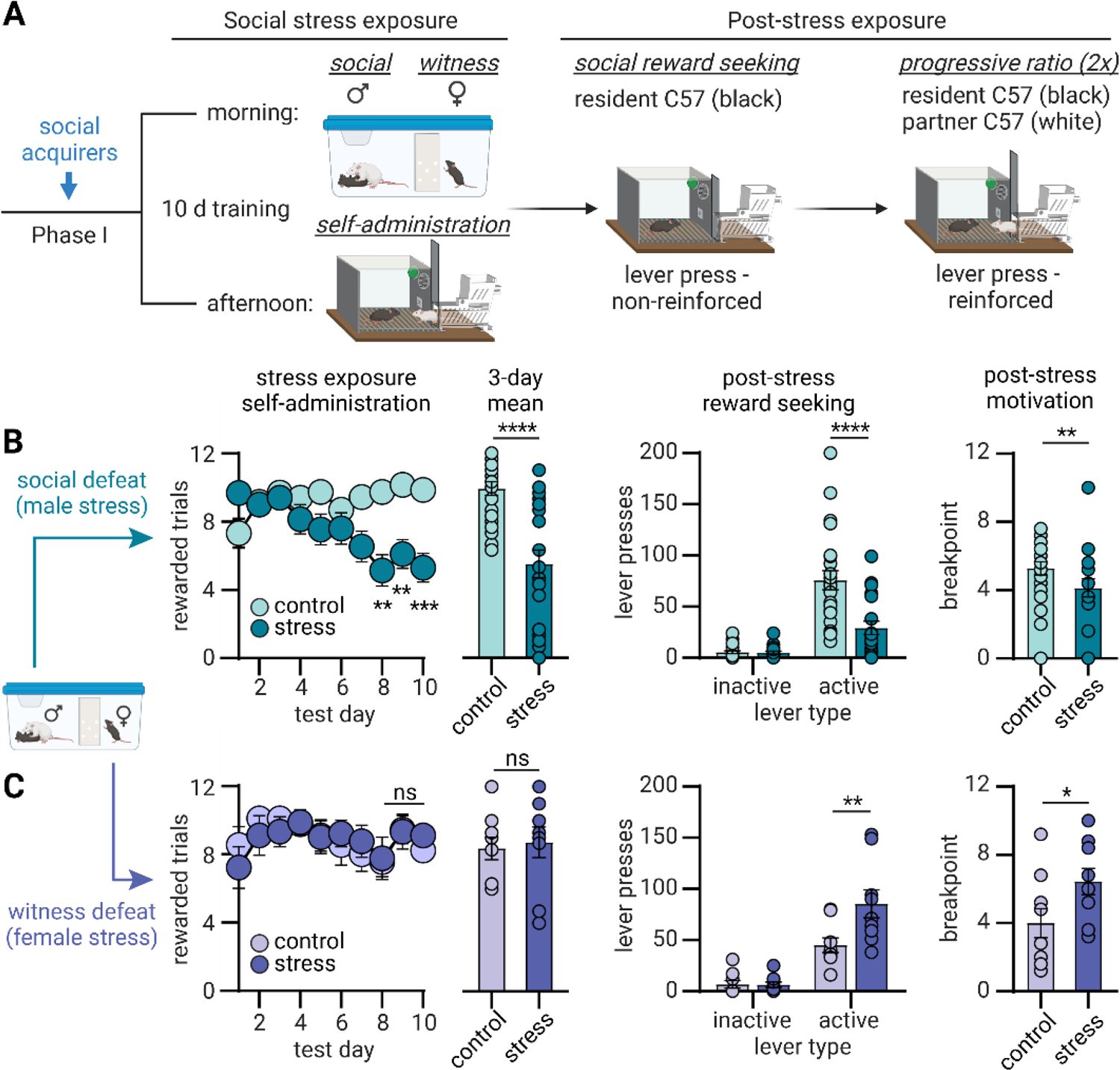
Social stress exposure impacts social self-administration and social reward seeking. (**A**) Experimental schematic for stress and post-stress phases of OSS. Male and female mice that acquired social self-administration were subjected to 10-d of once-daily social stress exposure followed 4-hr later by social self-administration test sessions. After the last day of social stress exposure, mice were tested for non-reinforced social reward seeking and 2-d of reinforced progressive ratio testing. (**B**) Male mice. <u>Left</u>: Number of rewarded trials over 10 days (12 trials/day) of social self-administration testing with male control (n=24) and stress exposed (n=20) mice, and mean rewarded trials from the final 3-d of social self-administration testing. <u>Middle</u>: Number of non-reinforced active and inactive lever presses during a 1-hr social reward seeking test under extinction conditions following social defeat stress exposure. <u>Right</u>: Mean breakpoint earned during two 2-hour progressive-ratio tests for social reward. (**C**) Female mice. <u>Left</u>: Number of rewarded trials over 10 days (12 trials/day) of social self-administration testing with female control (n=9) and stress exposed (n=9) mice, and mean rewarded trials from the final 3-d of social self-administration testing. <u>Middle</u>: Number of non-reinforced active and inactive lever presses during a 1-hr social reward seeking test under extinction conditions following social defeat stress exposure. <u>Right</u>: Mean breakpoint earned during two 2-hour progressive ratio tests for social reward. Individual data denoted with symbols. *p < .05. Data are mean ± SEM.

**Figure 4.**
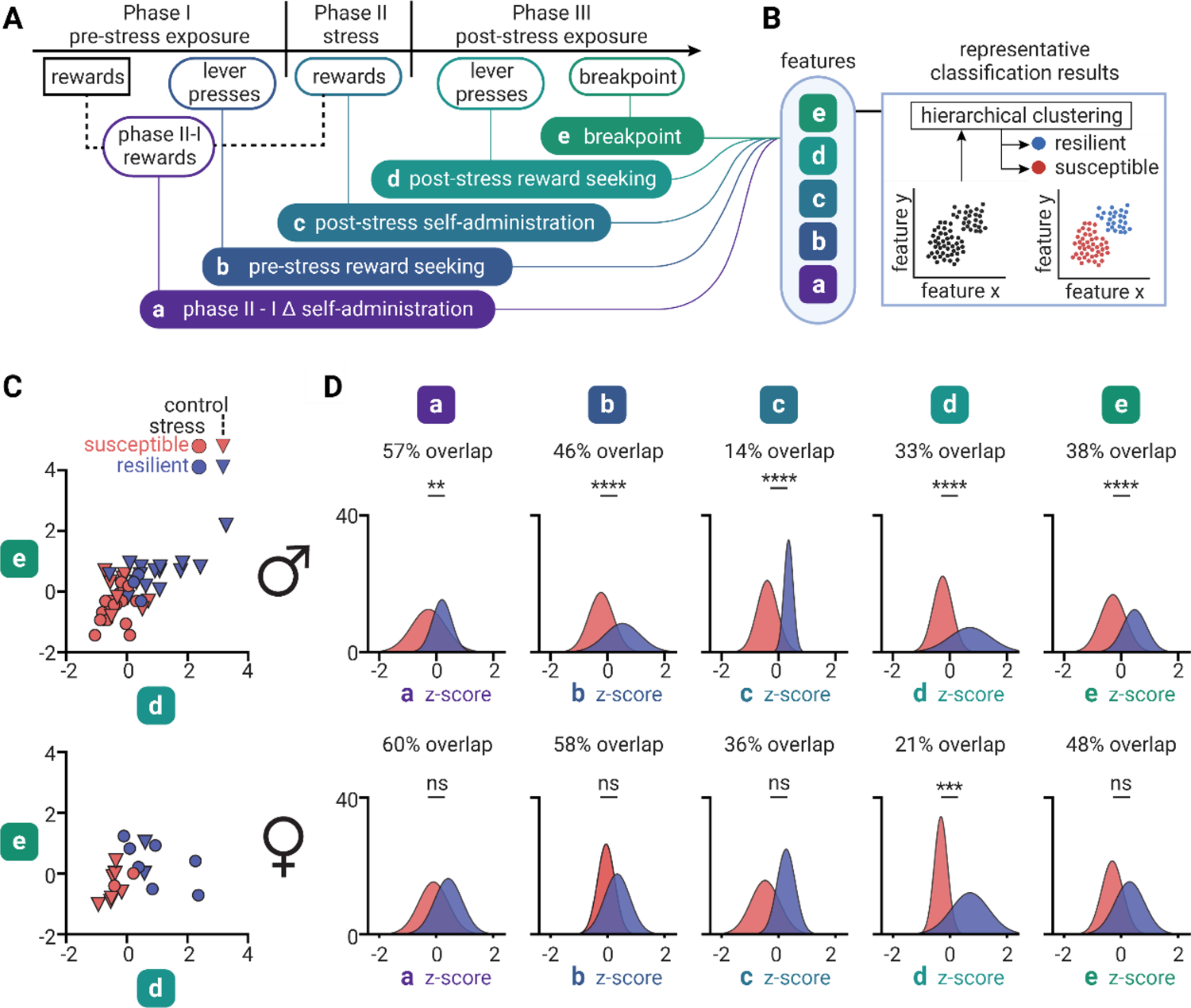
Social stress-induced hierarchical clustering of social reward phenotypes. (**A**) Analytical pipeline for hierarchical clustering of social operant metrics. Five operant metrics obtained during OSS served as features for clustering analysis: **a**, the difference in 3-day mean of social self-administration before and during stress (“*phases II - I Δ self-administration*”); **b**, lever pressing on first lever test (“*pre-stress reward seeking*”); **c**, the 3-day mean rewards obtained during social stress exposure (“*post-stress self-administration*”); **d**, lever pressing on second lever test (“*post-stress reward seeking*”); and **e**, mean breakpoint across the 2 days of progressive ratio testing (“*breakpoint*”). The five metrics were z-scored across experimental groups separately for male and female mice. (**B**) Schematic of unsupervised classification approach. Inset, representative hypothetical results for classification of social behavior during OSS. Clearly defined populations among operant data from OSS metrics (‘black dots’) should yield two discrete clusters, representing ‘susceptible’ (red) and ‘resilient’ (blue) populations of mice. (**C**) Male (top, n=44) and female (bottom, n=18) scatterplots of z-scored post-stress metrics (reward seeking and breakpoint) labeled by cluster assignment and stress condition. (**D**) Distribution of z-scored features (a-e) by cluster assignment for male (top, n=18 ‘resilient’) and female (bottom, n=9 ‘resilient’) mice. *p < .05.

### Exp. 1: Effect of social stress exposure on social self-administration and social reward seeking

The goal of Experiment 1 was to determine the consequence of social stress exposure on voluntary social self-administration and social reward seeking in male and female mice. In Phase I we trained 72 male and 25 female mice for self-administration and excluded 5 males and 1 female for inappropriate aggressive behavior or health concerns. 44 males and 18 females acquired social self-administration, defined as an average of 2 or more social rewards across the last 3 days of training (days 8-10). All mice were tested for non-reinforced social reward seeking (1-hr) the day following social self-administration training. During the social reward seeking tests lever presses led to contingent delivery of only the discrete cue previously paired with the delivery of the social partner. In Phase II, mice that acquired self-administration were placed into two groups per sex: male social defeat stress (n=20) and non-stressed controls (n=24), or female witness defeat stress (n=9) and non-stressed controls (n=9). In Phase III, across sequential days, we tested all male (n=44) and female (n=18) mice for non-reinforced reward seeking (1-hr) and 2 consecutive reinforced progressive-ratio tests.

### Exp. 2: Effect of social partner coat color on social self-administration

The goal of Experiment 2 was to determine the consequence of social partner coat color on social self-administration, controlling for the use of a contrasting coat color that improves manual and automated behavioral annotation. We trained a cohort of male black-coated C57Bl/6J mice paired with familiar, same coat color partners (n=20) and a cohort of male black-coated C57Bl/6J mice paired with familiar, different-coat color partners (n=24). We included both acquiring and non-acquiring mice in our analysis. Generalization of social self-administration across coat color is important for application of both manual annotation and pose-estimation and machine-guided automated behavioral classification^34–37^.

### Exp. 3: Effect of social partner familiarity and prior experience on social self-administration

The goal of Experiment 3 was to determine the consequence of partner familiarity on the acquisition of social self-administration. We trained black-coated male C57Bl/6J mice to self-administer a non-familiar (n=23) black-coated C57Bl/6J social partners and compared their acquisition of self-administration with the black-coated males from Exp. 2 (n=20) that had familiar partners. We included both acquirer and non-acquirer mice in our analysis. To assess how prior social experience may modulate acquisition of social self-administration, we trained and tested male C57Bl/6J mice as described above (‘naïve’, n=20), and then flipped (‘experienced’, n=20, within-subject) the roles of these mice with their familiar partners during subsequent self-administration and reward seeking testing. Familiarity of social self-administration partners has pragmatic implications for the design of social experiments and influences social behavior in mice. Previous studies show that C57Bl/6J mice show a preference towards non-familiar, novel social conspecifics compared to familiar mice in a 3-chamber social interaction test^38^. Previous studies of social self-administration in mice used same-sex non-familiar juvenile or age-matched conspecifics^18–20^.

### Exp. 4: Social stress-induced hierarchical clustering of social reward phenotypes

The goal of Experiment 4 was to cluster stress-susceptible and resilient phenotypes derived from operant behavioral metrics during OSS using a hierarchical clustering approach^17, 39–44^. We used a standard clustering algorithm (Ward’s method) to perform agglomerative hierarchical clustering on the animals’ operant OSS metrics. Five features were selected based on their predicted relevance to each animal’s expression of social motivation, and the changes they undergo following stress exposure: 1) the difference in mean social rewards obtained over the last 3 days of training (Phase I) and stress exposure (Phase II), 2) active lever presses from the pre-stress social reward seeking test, 3) mean social rewards obtained from the last 3 days of self-administration testing during stress exposure, 4) active lever presses from the post-stress social reward seeking test, and 5) mean breakpoint from progressive ratio tests. The detailed analytical pipeline is described in the Supplementary Methods. The clustering results were compared with the Calinski-Harabasz criterion^45^ using the differences in OSS features for male and female clusters, the overlap coefficient between their fitted distributions, and Kolmogorov-Smirnov tests. Clustering analyses and overlap coefficient calculations were performed using custom Python scripts.

### Exp. 5: Social stress-induced indexing of social reward behavior

In Experiment 5 we employed a different analytical approach to examine the spectrum of social motivation behavior exhibited by animals following social stress, using the same five operant OSS features from Exp. 4. The detailed analytical pipeline is described in the Supplementary Methods and results. Briefly, the five features were z-scored across mice of each sex and summed together to compose a summarizing measurement for each mouse as their cumulative ‘social index’ score. We first distributed the social index of males and females based on their experimental condition (’control’, ‘stress’) to examine their similarity, as described by their overlap coefficient. We then compared the raw OSS metrics of each animal to their social index score using linear regressions, to study the relationships between them before, during and following social stress exposure. To determine which features were most impactful for males and females during the procedure, we used PCA (3-component) on the z-scored features that composed the cumulative score and visually labeled each animal by their cumulative score value, subsequently assessing the correlation of each feature to the resulting principal components (Pearson’s *r*). Data scaling, linear regressions, overlap coefficient calculations and PCA were performed using custom Python scripts.

## Results

### Social self-administration and reward seeking in male and female mice

In Phase I of OSS, we used a trial-based fixed ratio (FR1) reinforcement schedule to determine whether mice would learn to lever press for freely moving affiliative social interaction with a familiar partner (Figure 2A). We trained male (n=72) and female (n=25) C57Bl/6J mice to self-administer for a familiar social partner (age and sex-matched white-coated C57Bl/6J; see Supplemental Methods). We excluded 4 males and 1 female that exhibited aggression or had health-related concerns. Of the remaining mice, 44 male and 18 female mice acquired (termed ‘acquirers’) and 23 male and 6 female (termed ‘non-acquirers’) failed to acquire social self-administration as measured by the daily number of rewarded trials (Figure 2B, left F_1,88_ = 76.8, p < .0001), with no difference between males and females (F_1,88_ = 0.475, p = 0.49). Mice that acquired social self-administration exhibited a significantly higher average of rewarded trials (F_1,88_ = 130.7, p < .0001) across the last 3 days of training compared to non-acquirers regardless of sex (Figure 2B, right, F_1,88_ = 0.112, p = 0.74). In the social reward seeking test, both male (F_1,176_ = 14.3, p = .001) and female (F_1,176_ = 14.3, p = .0003) acquirers had significantly greater active lever presses than non-acquirers (Figure 2C). Overall, a majority of male (65%, n=44) and female (75%, n=24) mice were classified as social acquirers following self-administration training. Complete statistical results can be found in Supplemental Table S1.

### Effect of familiarity, prior experience, and coat color on social self-administration and reward seeking

To test if coat color would impact acquisition of social self-administration, we compared the self-administration of male ‘different’ (n=20) coat color familiar partners to ‘same’ (n=24) coat color familiar partners. There was no effect of coat color across training days (F_1,40_ = 2.71, p = 0.11), but a significant difference in 3-day mean rewards (F_1,40_ = 2.71, p = 0.023) between ‘same’ and ‘different’ coat color acquirers (Figure 2D, left). In a reward seeking test using ‘same’ coat color familiar partners, acquirers exhibited significantly increased active lever responding (F_1,80_ = 33.9, p = < .0001) compared to non-acquirers (Figure 2D, right). Overall, 70% of males with partners of the ‘same’ coat color acquired self-administration (n=20).

We also asked how familiarity with a social partner impacted acquisition of social self-administration in male mice by using either familiar (n=20) and non-familiar (n=23) social partners. Familiarity had no effect on social self-administration (Figure 2E; F_1,39_ = 0.94, p = 0.34). Lastly, we examined whether prior social experience as a social partner influences later acquisition of social self-administration. Both ‘naïve’ (n=20) and ‘experienced’ (n=20) male mice (Supplemental Figure S1B) acquired social self-administration (F_1,36_ = 37.6, p < .0001), with a significant difference in 3-day mean (F_1,36_ = 3.01, p = 0.003). Both naïve (F_1,72_ = 25.5, p = .0008) and experienced (F_1,72_ = 25.5, p < .0001) acquirers exhibited higher social reward seeking (Supplemental Figure S1C) compared to non-acquirers, with a significant difference in active lever presses between naïve and experienced acquirers (F_1,36_ = 1.12, p = 0.036).

### Social self-administration and reward seeking in male and female mice following social stress exposure

Following Phase I acquisition of social self-administration, in Phase II we assigned mice into either ‘stress’ or no-stress ‘control’ groups. We exposed male and female mice to social defeat stress (n=20 stress, n=24 control) or witness defeat stress (n=9 stress, n=9 control), respectively, for 10 d paired with daily social self-administration test sessions (Figure 3A, left) to assess the consequence of social stress exposure on volitional self-administration of a familiar social partner. Stress males (Figure 3B) exhibited a reduction in rewarded trials compared to controls across days (F_1,42_ = 6.08, p = 0.018), resulting in a significantly lower average over the last 3 days (F_1,42_ = 26.68, p < .0001) of social self-administration testing. Stress females (Figure 3C) exhibited no difference in social self-administration compared to control females (F_1,16_ = 0.004, p = 0.95) across days.

In Phase III (Figure 3A, right), all mice were tested with a 1-hr non-reinforced lever test and 2 consecutive days of progressive ratio (PR) testing to gauge social reward seeking and motivation, respectively, following social stress exposure. Stress males (Figure 3B) pressed the active lever significantly less (F_1,84_ = 12.1, p < .0001) during non-reinforced reward seeking, and had a significantly lower breakpoint (F_1,42_ = 8.46, p = 0.006), than control males. Stress females (Figure 3C) pressed the active lever significantly more (F_1,32_ = 6.43, p < .0001) during non-reinforced reward seeking, and exhibited a higher breakpoint, compared to control females (F_1,17_ = 4.46, p = 0.0499).

### Cluster analysis of stress susceptibility and resilience in operant social behavior exhibited across OSS

Next, we used the data to evaluate social motivation following social stress exposure (Figure 4A). Preclinical models of social stress-related behavior typically assign individual differences into distinct subpopulations that show stress susceptibility or resilience. The most common approach is to use variations on chronic social defeat stress, which introduce high throughput and standardization through forced social interactions, but then omit volitional social motivation metrics in their behavioral readout^26^. To determine whether individual differences in social self-administration and reward can be non-arbitrarily classified into susceptible and resilient classifications (Figure 4B), we used a hierarchical clustering algorithm (Ward’s method, see Supplemental Methods and Materials) to perform an unsupervised analysis of OSS metrics as features to generate two behavioral clusters (‘resilient’ and ‘susceptible’). We selected five OSS *features*: **a**, the difference in 3-day mean of social self-administration before and during stress (“phases II - I Δ self-administration”); **b**, lever pressing on first lever test (“pre-stress reward seeking”); **c**, the 3-day mean rewards obtained during social stress (“post-stress self-administration”); **d**, lever pressing on second lever test (“post-stress reward seeking”); and **e**, mean breakpoint across progressive ratio testing (“breakpoint”).

To describe these clusters in relation to the final OSS behavior, we compared the two post-stress tests (post-stress reward seeking and breakpoint values) for male (Figure 4C, top) and female (Figure 4C, bottom) mice that were putatively classified into the ‘susceptible’ (male n= 26, female n= 9) or ‘resilient’ (male n= 18, female n= 9) clusters. 17 stress males and 2 stress females were classified as ‘susceptible’, and 3 stress males and 7 stress females as ‘resilient’. 9 control males and 7 control females were classified as ‘susceptible’, and 15 control males and 2 control females as ‘resilient’. As a result, the males had a 27.3% overlap in classification between stress and control groups, while the females had a 77.8% overlap. Moreover, male clusters were more discrete than female clusters based on Calinski-Harabasz scores of 26.4 and 9.26, respectively. The distribution of OSS features grouped by cluster assignment shows that for males (Figure 4D, top) the most discriminating features were *Δ self-administration*, *post-stress self-administration* and *breakpoint* (Kolmogorov-Smirnov; **a**, 57% overlap, p = 0.001; **d**, 33% overlap, p < .0001; **e**, 38% overlap, p < .0001), and female (Figure 4D, bottom) clusters were separated the most by *post-stress reward seeking* (**d**, 21% overlap, p = 0.0007). These results suggest that the behavioral spectrum of social self-administration and reward in male and female mice following social stress exposure is not clearly defined by two clusters, indicating that a different approach is required to describe this relationship.

### Individual differences in social motivation following social stress exposure during OSS

Our clustering results (Figure 4C-D) suggested that OSS features capture varying facets of social motivation before, during and after social stress. The differences between stress mice compared to their non-stress controls (Fig. 3B-C), and the variability observed within mice among features, suggest that, rather than using categorical classification, OSS individual variability is better modeled as a distribution along an integrative spectrum. To test this hypothesis, we aggregated the five OSS features, combining them into a cumulative score for each animal (termed ‘social index’; Figure 5A) that represents a summary of their social reward and motivation behavior. Consistent with previous behavioral results, stress males (Figure 5B, left) occupied the lower end of the social index distribution (32% overlap; Kolmogorov-Smirnov, p < .0001), while the distribution for stress females (Figure 5B, right) was higher than control females (68% overlap, p = 0.126).

**Figure 5.**
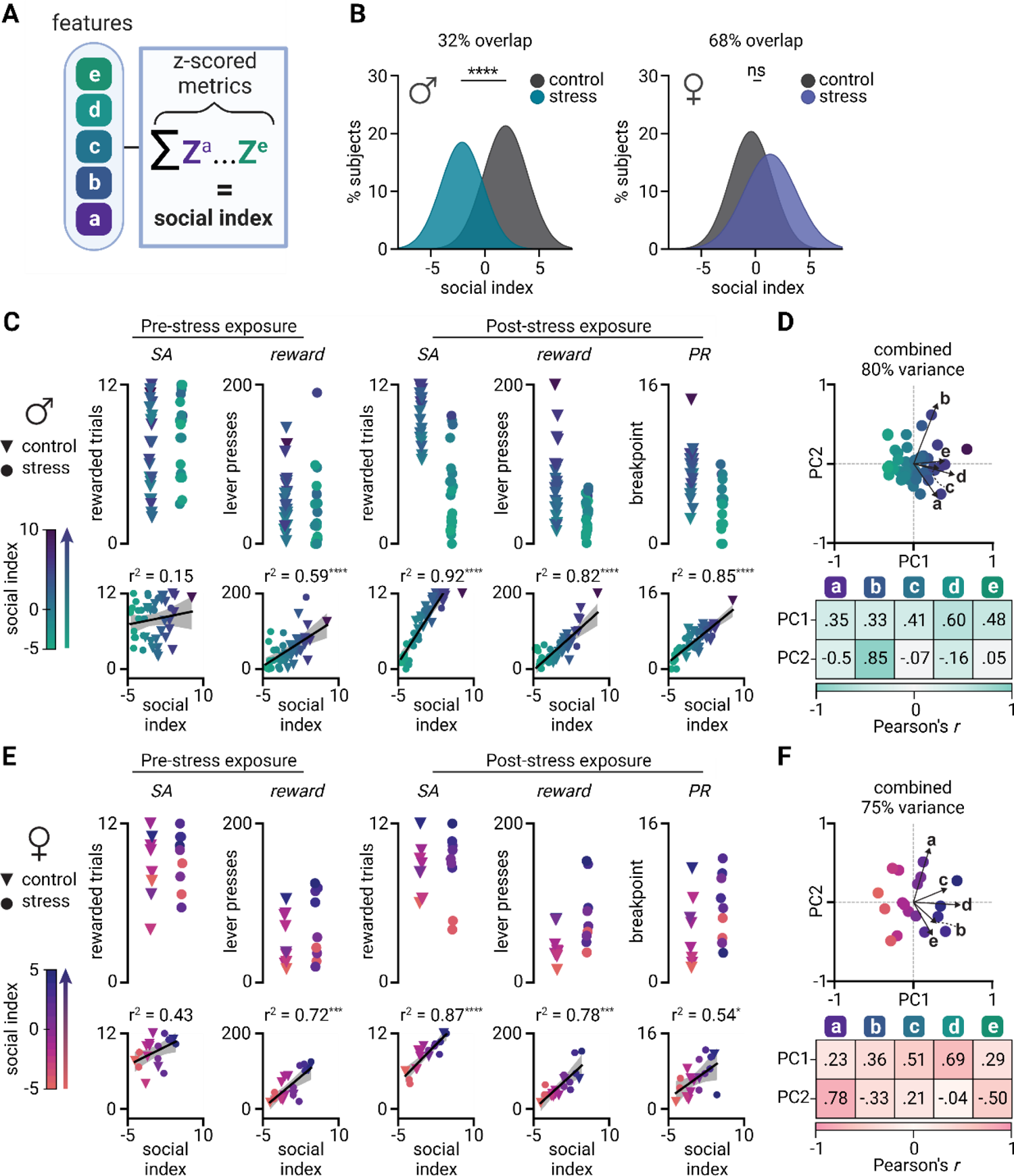
Social stress-modulated indexing of social reward behavior. (**A**) Social index calculation. The five z-scored metrics chosen (Z^a^-Z^e^) for clustering analysis were aggregated per animal to obtain a cumulative sum of their features (‘social index’). (**B**) Social index distributions for males (left, n=44) and females (right, n=18). (**C**) <u>Top</u>: Raw OSS metrics for control and stress males, color-mapped by their social index score. <u>Bottom</u>: Correlation of social index with individual features. (**D**) Biplot of PCA for features from males. (**E**) <u>Top</u>: Raw OSS metrics for control and stress females, color-mapped by their social index score. <u>Bottom</u>: Correlation of social index with individual features. (**F**) Biplot of PCA for features from females. *SA: self-administration*. *p < .05.

We next explored how each OSS feature related to the social index distributions. For males (Figure 5C), four of the five OSS metrics correlated with each animal’s social index score (*pre-stress reward seeking*: *r_2_* = .59, p < .0001; *post-stress self-administration*: *r_2_* = .92, p = < .0001; *post-stress reward seeking*: *r_2_* = .82, p < .0001; *PR*: *r_2_* = .85, p < .0001). We used principal component analysis (PCA) to explain the feature-dependent variability across animals. In males (Figure 5D) the first principal component (PC) that best described their social index distribution (60% variance) was most positively correlated with post-stress reward seeking (**d**, *r* = 0.60) and breakpoint (**e**, *r* = 0.48), while the second PC (20% variance) was positively correlated with pre-stress reward seeking (**b**, *r* = 0.85) and had negative correlations with Δ self-administration (**a**, *r* = −0.5) and post-stress reward seeking (**d**, *r* = −0.16). For females (Figure 5E), four out of five OSS metrics correlated with each animal’s social index score (*pre-stress reward seeking*: *r_2_* = .72, p < .001; *post-stress self-administration*: *r_2_* = .87, p = < .0001; *post-stress reward seeking*: *r_2_* = .78, p < .0001; *PR*: *r_2_* = .54, p = 0.02). In females (Figure 5F) the first PC (50% variance) correlated strongly with post-stress reward seeking (**d**, *r* = 0.69) and post-stress self-administration (**c**, *r* = 0.51), while the second (25% variance) was positively correlated with Δ self-administration (**a**, *r* = 0.78) and negatively correlated with breakpoint (**e**, *r* = −0.5). These results further demonstrate that OSS can be used to detect changes in social behavior due to stress that exist across multiple dimensions of operant behavior.

### Stability of the social index

Next, we asked how well operant social behavior from earlier OSS phases could predict final social index scores. Accurate stratification of individual social index positions using early operant features would be beneficial for future experimental applications. We compared the individual social index score distributions and their ranked order (Figure 6A) using 4 and 3 OSS features (without PR or both PR and breakpoint, respectively) to those derived from all 5 features. Figure 6B-C (top) show the distributions of social index scores calculated for control (‘triangle’) and stress (’circle’) males and females as features are reduced. To measure and test the strength of the relationships between the three feature spaces and their resulting indices, we performed multiple linear regression analyses on each and compared these regressions. Reduced feature spaces for data from males (Figure 6B, bottom) had similar correlations between predicted and actual social index scores to that of the full feature space (‘5 features’, *r_2_* = 1, p = < .0001; ‘4 features’, *r_2_* = .98, p = < .0001; ‘3 features’, *r_2_* = .89, p = < .0001). Analysis of covariance (ANCOVA) on the regression models for each feature space showed significant differences between the slopes (p = 0.021), indicating that reduced feature spaces may approximate the final index score distribution. For females (Figure 6C, bottom), the regressions of reduced feature sets were not significantly different from using all 5 features (‘5 features’, r_2_ = 1, p = < .0001; ‘4 features’, r_2_ = .94, p = < .0001; ‘3 features’, r_2_ = .89, p = < .0001; slopes, p = 0.36; intercepts, p > 0.99). Together, these results suggest that removal of Phase III OSS metrics may lead to slightly different predictions for social index score distributions of males. However, the differences between social index score distributions are highly correlated with their respective feature set.

**Figure 6.**
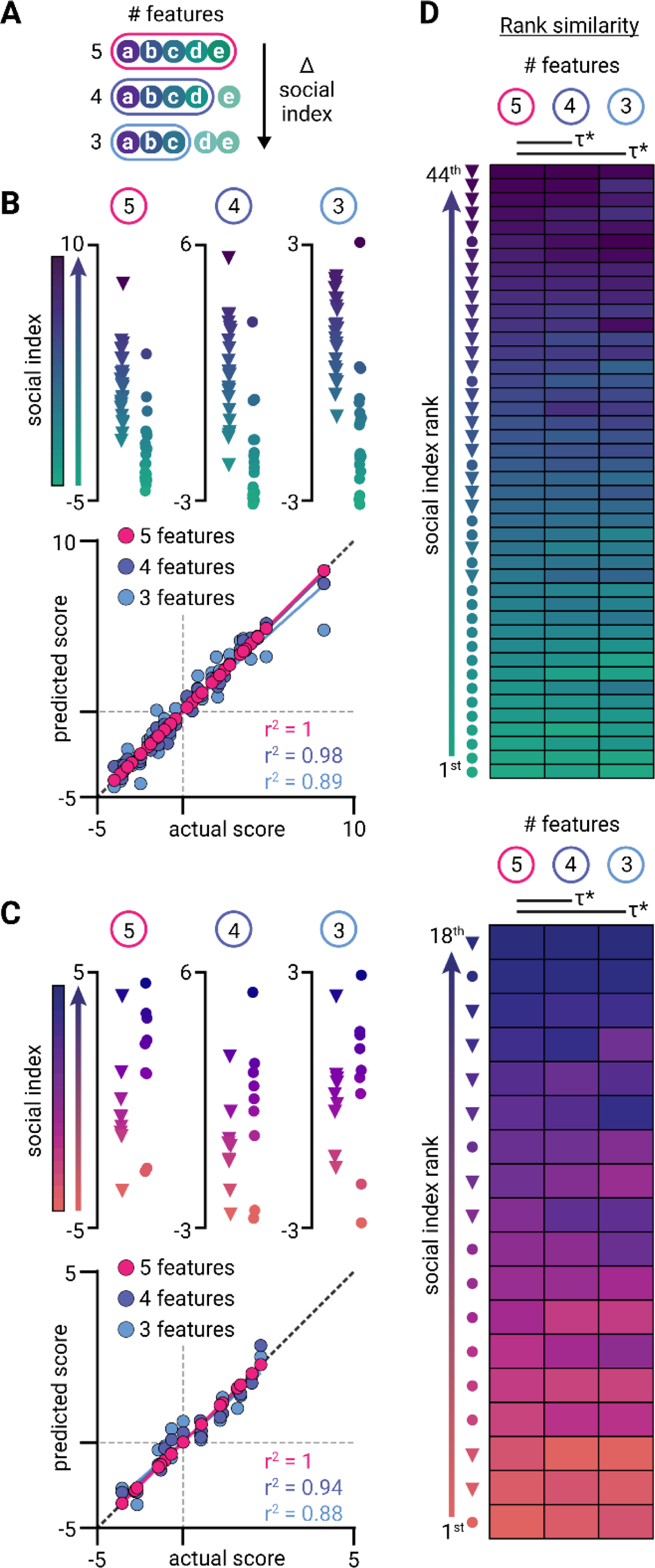
Stability of social index ranking. (**A**) Stability test schematic. To determine how well earlier features can predict the outcome of OSS, the social index for each animal was calculated without PR (‘4 features’, purple) and without PR or post-stress reward seeking (‘3 features’, light blue), and compared against the social index obtained using all 5 features (pink). (**B**) <u>Top</u>: Male distributions of social index scores using 5, 4, or 3 OSS features. <u>Bottom</u>: Male multiple linear regression-predicted scores using 5, 4, or 3 OSS features correlated against the 5 feature OSS social index distribution. (**C**) <u>Top</u>: Female distributions of social index scores using 5, 4, or 3 OSS features. <u>Bottom</u>: Female multiple linear regression-predicted scores using 5, 4, or 3 OSS features correlated against the 5-feature OSS social index distribution. (**D**) Male (top) and female (bottom) heatmaps depicting the ordered ranking of social index scores (rows; controls, triangles; stress, circles) as a function of OSS features (columns). τ: Kendall’s tau; *p < .05.

We next tested the stability of individual social index rankings as a function of decreasing OSS feature space, by ranking the social index scores for each feature set for males and females (Figure 6D, top and bottom, respectively). The ranked order of social index scores for control (triangle) and stress (circle) mice, from lowest to highest social index, is shown in mice ordered by their full (5 feature) social index ranking. Rank similarity (Kendall’s tau) tests found the rankings of social index scores from reduced feature spaces to be significantly similar to the final ranking using all features for both male (5 v 4 features, tau = 0.90, p = < .0001; 5 v 3 features, tau = 0.82, p < .0001) and female (Fig. 6D, bottom; 5 v 4: tau = 0.91, p < .0001; 5 v 3: tau = 0.80, p < .0001) mice. These results show the stability of social index score distributions.

## Discussion

We used volitional social self-administration procedures to evaluate social motivation following social stress exposure in male and female mice. We named this procedure operant social stress (OSS), which is composed of three phases: (*i*) social self-administration training, (*ii*) social stress exposure concurrent with daily reinforced self-administration to test social motivation, and (*iii*) operant social reward seeking under both non-reinforced and reinforced conditions. Male and female mice similarly acquired social self-administration for familiar age- and sex-matched social partners using a trial-based design that was reinforced with freely moving social interaction. Following acquisition of social self-administration, male and female mice exhibited similar social reward seeking under non-reinforced extinction conditions. Changing the familiarity or coat color of the social partner did not modify acquisition of social self-administration.

Social stress exposure during OSS had a lasting impact on social motivation in both male and female mice. Compared to non-stressed male controls, male mice exposed to social defeat exhibited a decrease in social self-administration across defeat days followed by decreased social reward seeking and motivation in both non-reinforced and reinforced conditions. Compared to non-stressed female controls, female mice exposed to witness defeat maintained similar levels of social self-administration followed by increased social reward and motivation. Using standard hierarchal clustering approaches based on operant OSS metrics, we explored individual variability in social motivation and observed that binary classification (i.e., susceptible or resilient) does not adequately describe variability in either male or female mice. Rather, individual variability in social motivation following stress was better described using a ‘social index’ score derived from the sum of z-scored operant OSS features. We found that the social index distributions of individual mice are stable regardless of the number of OSS features used in the derivation. Overall, our data indicate that the inclusion of volitional social self-administration effectively surveys the spectrum of stress-induced individual variability in social motivation. The OSS procedure thus provides a platform for the study of neural mechanisms driving changes in volitional social motivation in both male and female mice.

### Inbred male and female mice acquire social self-administration and exhibit social reward seeking

Both male and female inbred C57Bl/6J mice acquired social self-administration using a trial-based procedure that is reinforced by freely moving physical social interaction and exhibited social reward seeking under non-reinforced extinction conditions. We have previously used a similar social self-administration procedure in male CD-1^14, 17^ and female CFW^16^ outbred mice to examine behavioral, cell-type and circuit-specific mechanisms of aggression reward seeking. Similar operant procedures have been used to study aggression reward for nearly two decades (see Golden et al., 2019 for a comprehensive review of aggression social-self administration procedures)^15^.

More recent work has adapted operant aggression self-administration procedures to assess operant affiliative social reward, using barrier-based sensory contact procedures where operant actions are reinforced by sensory access to a social partner that is behind a perforated barrier. These studies have shown that male C57Bl/6J mice will self-administer sensory access to juvenile male social partners^18, 20, 46^, and that outbred female CD-1 but not inbred female C57Bl/6J mice will self-administer sensory access to age-matched social partners^19^. Our data are the first to show that both male and female inbred C57Bl/6J mice will self-administer familiar age-matched affiliative social partners when reinforced with freely moving social interaction. This may be because aspects of physical social interaction are more rewarding than sensory contact in mice, but such direct comparisons have yet to be published and should be an area of future investigations. This is supported by studies in mice examining the rewarding nature of social touch^47^ under involuntary social pairing during allogrooming, and similar mechanisms may modulate or contribute to volitional social reward seeking.

Notably, there are inherent pragmatic trade-offs to using sensory contact versus physical contact social self-administration procedures. Due to the absence of physical contact and the subsequent necessity to physically separate social partners, sensory contact procedures are streamlined for high-throughput and minimal experimenter involvement. However, that same absence creates interpretive difficulties in understanding the resident’s motivation for performing the operant action. In the present manuscript, several male mice were excluded during social self-administration training for displaying aggressive, rather than affiliative, social behaviors during self-administration training. Without observation of these physical interactions, their inclusion may have led to erroneous interpretation of affiliative social motivation.

### Stress-induced modulation of volitional social motivation in male and female mice

Stress resilience and susceptibility in rodents are associated with neurophysiological changes in the mesolimbic dopaminergic system^48–53^. However, this mechanistic understanding is constrained by two pragmatic procedural considerations. First, variations on social stress models like chronic social defeat stress and witness defeat stress allow for high-throughput and standardized procedures but use involuntary social interactions as the key metric of stress susceptibility or resilience. This removes the ability to ask fundamental questions about how the brain encodes and controls volitional social motivation either during or after stress exposure. Second, since these procedures were initially developed to generate large cohorts of stress susceptible and resilient mice for molecular and genomic analysis, they stratify mice based on the single metric of passive social exploration during a rapid social interaction test. This fails to capture the multi-dimensionality of individual variability in responses to stress.

To better understand reward-related social behavior, there is a need to incorporate volitional social motivation within the context of social stress. The OSS procedure helps to fill this gap by providing a social self-administration procedure that may be repeated at multiple time points during and after social stress exposure to evaluate changes in voluntary social reward-seeking behavior. Our findings in male mice correspond conceptually with previous observations using social interaction testing, where susceptible mice show social avoidance, as stressed male mice attenuated social self-administration of familiar social partners across social defeat exposures compared to non-stressed controls. Similarly, using both non-reinforced social reward seeking tests and reinforced progressive ratio tests, stressed males exhibited decreased social motivation compared to non-stressed controls. These results replicate and extend findings from previous models, showing that anxiety-like behaviors due to social defeat stress, such as social avoidance, are additionally expressed in the volitional social behavior of male mice as a reduction in motivation for affiliative interactions with familiar social partners as social reward.

Conversely, females that were exposed to witness defeat stress exhibited similar social self-administration of familiar social partners throughout the social stress exposure, and elevated social reward seeking and motivation after stress exposure, compared to control females. These data are reminiscent of previously reported witness defeat results in females that have found signs of heightened arousal, indicated by anxiogenic behavior, corticosterone levels in serum, proinflammatory markers, among others, which are suggested to describe a hypervigilance state^27, 54, 55^. Such hypervigilance has also been reported in females undergoing social defeat stress^56^. However, our witness defeat-stressed social self-administration results are different than previous reports of social avoidance^54^, which may be a result of the involuntary versus voluntary nature of the tested social interactions.

Importantly, our current data does not provide a direct comparison of sex differences between male and female mice, as the social stress exposures were different. Rather, here we take advantage of the OSS procedure to compare stressed mice to their same-sex controls, revealing unexpected differences in social decision making and reward seeking between sexes. We designed OSS to be modular, and future iterations will incorporate variations of female social defeat stress^54, 56–60^ and male witness defeat stress^27^, to allow direct behavioral and neurophysiological comparisons across sexes.

### Individual variability in social reward seeking following stress

We demonstrate that multidimensional operant features allow for the identification of individual differences in volitional social motivation following stress, and further incorporate volitional social behavior as an endpoint for the highly used social defeat and witness defeat stress models in male and female mice, respectively. We have previously used cluster analysis to identify discrete subpopulations of aggression reward seeking mice using operant procedures^17^, and we applied a similar approach using OSS. Contrary to these earlier findings, our hierarchical clustering analysis on a subset of operant features during OSS did not accurately describe individual differences in social reward seeking following stress. Rather, this analysis yielded clusters that discerned distributions in some features but not others, ultimately failing to describe the breadth of behavioral differences we observed in both male and female mice.

To describe this spectrum of individual variability we used a method with minimal data transformation to provide a descriptive summary of each animal’s social decision-making and reward seeking behavior, while preserving the diversity of behavioral expression found across features^61^. We aggregated the key operant features from OSS into a social index score for each animal that contained a summary of their social operant behaviors. We show that the social index score relates back to its individual OSS components, and that the distribution of social indices of male and female mice accurately describes the spectrum of individual variability. A benefit of OSS experimental design is the longitudinal tracking of operant responses before, concurrent with, and after social stress exposure. This design revealed the absence of a relationship between the acquisition of social self-administration training and social index score, and rather the importance of social stress exposure during OSS leading to meaningful differences in social decision-making and reward behavior. Notably, we show that the social index score is stable even with the exclusion of operant features following stress exposure, providing flexibility in the use of the OSS procedure in combination with modern neural recording and manipulation approaches.

### Technical advances supported by OSS

In addition to the conceptual advancement OSS provides by incorporating volitional social behavior within the context of social stress exposure, we designed OSS with compatibility of emerging neuroscience methods in mind. First, we validated the use of differently coat colored social partners during social self-administration to facilitate the future use of multi-animal pose-estimation and machine-guided automated behavioral classification. Although significant advances^35–37^ have reduced unintended ID swapping, the annotation of robust and rapid freely moving social interaction is simplified when the subjects are visually distinguishable. The addition of both supervised^34^ and unsupervised^62–64^ behavioral classification within the context of volitional social decision making has tremendous potential for resolving underlying neural mechanisms. Second, since OSS incorporates time-locked operant events, as well as discriminative and conditioned cues, physiological measurements and neural manipulations can be paired with discrete events within the OSS model to study neural circuitry during social reward behavior^65^.

## Conclusion

We extend the use of social defeat stress and witness defeat stress to include operant metrics of social motivation and introduce the social index as an approach to describe individual differences. These data conceptually align with the social processes domain of the Research Domains Criteria (RDoC) system^66^ that states pathology presents as a spectrum; this spectrum is reflected in the social index distributions we observed. Overall, we provide a new procedure that may be used for the identification of neural mechanisms underlying individual differences and changes in social motivation following social stress exposure.

## Figure Legends

**Supplementary Figure 1.**
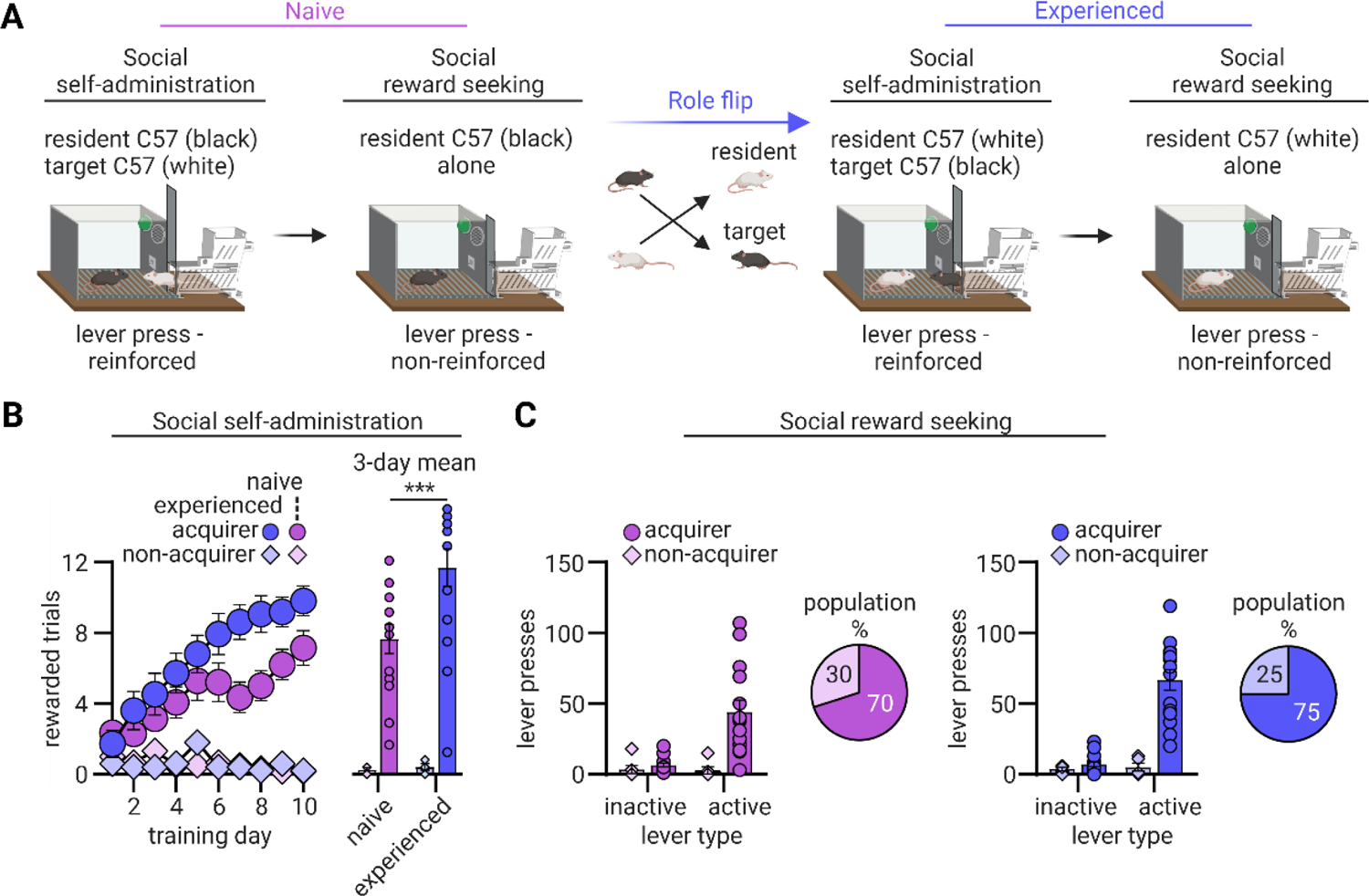
*Prior experience on social self-administration.* (**A**) Prior experience experiment schematic. To test how prior experience may influence social self-administration acquisition, a cohort was first trained and tested as described above (left, ‘Naïve’), after which the roles were flipped with their conspecifics, who were subsequently trained and tested as well (‘Experienced’). (**B**) <u>Left</u>: Number of rewarded trials over 10 days (48-minute session/day; 12 trials/day) of social self-administration under a trial-based fixed-ratio 1 reinforcement schedule with naïve (n=20) and experienced (n=20) resident male mice. <u>Right</u>: Mean rewarded trials from the final 3-d of social self-administration. (**C**) Number of non-reinforced naïve (left) and experienced (right) active and inactive lever presses during a 1-hr social reward seeking test under extinction conditions following social self-administration. Pie charts show the proportion of each cohort that acquired social self-administration. *p < .05. Data are mean +/- SEM.

## Supplementary Methods

We describe the specific experiments in the main text methods and provide the detailed description of experimental subjects, apparatus, and general procedures here.

### The operant social stress (OSS) procedure

Operant social stress (OSS) is composed of three phases: (*i*) social self-administration training and reward seeking, (*ii*) social stress exposure concurrent with social self-administration testing, and (*iii*) post-stress operant social reward testing under both non-reinforced and reinforced conditions (Figure 1, see “General operant self-administration experimental procedures” below for detailed descriptions). In Phase I, experimental male and female black-coated C57BL/6J mice are trained to press a lever for freely moving physical access to a familiar same-sex conspecific partner (white-coated C57BL/6J for compatibility with behavioral annotation) in a trial-based, 12-trial/day design for 10 days (Figure 1A and 2A). Following social self-administration acquisition, social reward seeking is measured using a 1-hr non-reinforced test under extinction conditions. In Phase II, male mice are exposed to social defeat stress and female mice to witness defeat stress, once-daily for 10 days, and 4 hr after each daily stress exposure they undergo social self-administration testing with their social partner (Figure 1B and 3A, left). Lastly, in Phase III, experimental mice are tested for non-reinforced social reward seeking and reinforced social progressive ratio responding (Figure 1C and 3A, right). These operant metrics are used to stratify mice according to social stress-induced dysregulation of social self-administration and reward seeking (Figures 4-6).

### Exp. 1: Effect of social stress exposure on social self-administration and social reward seeking

The goal of Experiment 1 was to determine the consequence of social stress exposure on voluntary social self-administration and social reward seeking in male and female mice. In Phase I we trained 72 male and 25 female mice for self-administration and excluded 5 males and 1 female for inappropriate aggressive behavior or health concerns. 44 males and 18 females acquired social self-administration, defined as an average of 2 or more social rewards across the last 3 days of training (days 8-10). All mice were tested for non-reinforced social reward seeking (1-hr) the day following social self-administration training. During the social reward seeking tests lever presses led to contingent delivery of only the discrete cue previously paired with the delivery of their social partner. In Phase II, mice that acquired self-administration were placed into two groups per sex: male social defeat stress (n=20) and non-stressed controls (n=24), or female witness defeat stress (n=9) and non-stressed controls (n=9). In Phase III, across sequential days, we tested all male (n=44) and female (n=18) mice for non-reinforced reward seeking (1-hr) and 2 consecutive reinforced progressive-ratio tests.

### Exp. 2: Effect of social partner coat color on social self-administration

The goal of Experiment 2 was to determine the consequence of social partner coat color on social self-administration, controlling for the use of differently coated partners which improves manual and automated behavioral annotation. We trained a cohort of male black-coated C57Bl/6J mice paired with familiar same color coat partners (n=20) and a cohort of male black-coated C57Bl/6J mice paired with familiar different coat color partners (n=24). We included both acquiring and non-acquiring mice in our analysis. Generalization of social self-administration across coat color is important for easier use with both manual annotation and with pose-estimation and machine-guided automated behavioral classification^34–37^.

### Exp. 3: Effect of social partner familiarity and prior experience on social self-administration

The goal of Experiment 3 was to determine the consequence of partner familiarity on the acquisition of social self-administration. We trained black-coated male C57Bl/6J mice to self-administer a non-familiar (n=23) black-coated C57Bl/6J social partners and compared their acquisition of self-administration with the black-coated males from Exp. 2 (n=20) that had familiar partners. We included both acquirer and non-acquirer mice in our analysis. To assess how prior social experience may modulate acquisition of social self-administration, we trained and tested male C57Bl/6J mice as described above (‘naïve’, n=20), and then flipped (‘experienced’, n=20, within-subject) the roles of these mice with their familiar partners during subsequent self-administration and reward seeking testing. Familiarity of social self-administration partners has pragmatic implications for the design of social experiments and has been found to influence social behavior in mice. Previous studies show that C57Bl/6J mice show a preference towards non-familiar novel social conspecifics compared to familiar mice in a 3-chamber social interaction test^38^. Subsequent studies of social self-administration also examine same-sex non-familiar juvenile or age-matched conspecifics^18–20^.

### Exp. 4: Social stress-induced hierarchical clustering of social reward phenotypes

The goal of Experiment 4 was to cluster stress-susceptible and resilient phenotypes derived from operant behavior during OSS using standard hierarchical clustering approaches. We used a standard clustering algorithm (Ward’s method) to perform agglomerative hierarchical clustering on the animals’ operant OSS metrics. These features were selected based on their predicted relevance to each animal’s expression of social decision-making and motivation, and the changes they undergo following stress exposure: 1) the change in mean social rewards obtained over the last 3 days of training (Phase I) and stress exposure (Phase II), 2) active lever presses from the pre-stress social reward seeking test, 3) mean social rewards obtained from the last 3 days of self-administration testing during stress exposure, 4) active lever presses from the post-stress social reward seeking test, and 5) mean breakpoint from progressive ratio tests. The clustering results were compared using the Calinski-Harabasz criterion^45^, using the differences in OSS features for male and female clusters, the overlap coefficient between their fitted distributions, and Kolmogorov-Smirnov tests. Clustering analyses and overlap coefficient calculations were performed using custom Python scripts.

### Exp. 5: Social stress-induced indexing of social reward behavior

In Experiment 5 we employed a different analytical approach to examine the spectrum of social motivation behavior exhibited by animals following social stress, using the same operant OSS features from Exp. 4. Briefly, the five metrics were z-scored across mice of each sex and summed together to compose a summarizing measurement for each mouse as their cumulative ‘social index’ score. We first distributed the social index of males and females based on their experimental condition (’control’, ‘stress’) to examine their similarity, as described by their overlap coefficient. We then compared the raw OSS metrics of each animal to their social index score using linear regressions, to study the relationships between them before, during and following social stress exposure. To determine which features were most impactful for males and females during the procedure, we used PCA (3-component) on the z-scored features that composed the cumulative score and visually labeled each animal by their cumulative score value, subsequently assessing the correlation of each feature to the resulting principal components (Pearson’s *r*). Data scaling, linear regressions, overlap coefficient calculations and PCA were performed using custom Python scripts.

### Subjects

Our experimental subjects were ∼20-25 g 8-12-week-old sexually naïve male and female C57BL/6J mice (bred in-house from Jackson Lab stock #000664). For social partners during operant social self-administration, we used age and size-matched male and female 8-12-week-old white-coated C57BL/6J mice (B6Cg-Tyr^c-2J^/J, bred in house from Jackson Lab stock #000058, n=72 male, n=25 female), of which 6 were excluded from analysis (5 males). We chose to use white-coated social partners to facilitate subject identification, which is improved by using social pairs with different coat colors. For control experiments, to examine how coat color (n=44), familiarity (n=43) and prior experience (n=40) influence social self-administration, we used standard black-coated age and sex-matched C57BL/6J mice (bred in-house from Jackson Lab stock #000664). For screening aggressive CD-1 residents prior to social defeat and witness stress we used ∼20-25 g 8-12-week-old sexually naïve male C57BL/6J male mice (bred in-house from Jackson Lab stock #000664, n=20), due to their well-established ethological characterization as subordinate to CD-1 mice in chronic social defeat stress^22, 26^.

For resident mice in social defeat stress, we used ∼40 g 4-6-month-old sexually experienced male CD-1 mice (Charles River Labs, CRL, n=30). We confirmed with CRL animal-facility staff that all the sexually experienced CD-1 males had equal access to receptive females. CRL’s procedure is to pair-house individual males with several females from PD28 until purchase. Pregnant females are switched with new non-pregnant females, with no break between cycles. Males that do not successfully breed are removed from the breeding pool and not made available for purchase.

We gave all mice free access to standard food chow and water in all experiments. We pair-housed all experimental C57Bl/6J mice with enrichment (cotton padding) in standard Allentown clear polycarbonate cages covered with stainless-steel wire lids at least one week prior to experiments, and we maintained them on a reverse 12-h light/dark cycle (light off at 0900 am). We group-housed the non-experimental male C57Bl/6J mice for CD-1 aggression screening. Aggressive CD-1 male mice were singly housed one week prior to social defeat screening. We performed all experiments in accordance with the Guide for the Care and Use of Laboratory Animals (8th edition; 2011), under protocols approved by the Animal Care and Use Committee.

## Apparatus

### Operant social self-administration apparatus

The operant social self-administration apparatus was adapted from previously published studies, schematically detailed in Fig. 2A. We trained and tested all mice in standard Med Associates operant chambers. Each chamber was enclosed in a ventilated sound-attenuating cubicle and illuminated by one of two houselights, each positioned above two retractable levers on opposite sides of the chamber. These two retractable levers were designated “active,” and a third non-retractable lever was designated “inactive”; all levers were positioned 2.4 cm above the grid floor. During social self-administration experiments, presses on the designated active lever (only extended during social self-administration) resulted in delivery of a same-sex conspecific partner mouse and 2-s tone cue (2900 Hz, 20 dB above background). For all social experiments, we presented the partner through an automatic guillotine-style door adjacent to the active lever.

To facilitate social conspecific presentation in all operant experiments, we attached a custom-made 3D-printed two-level mouse delivery device to each operant behavior chamber; the delivery device housed the social conspecific during social self-administration sessions. Upon completion of the reinforcement-schedule requirement for lever presses (FR1 or PR) and presentation of the conditioned tone cue, the automatic guillotine door opened vertically for 10-s, and we guided the social partner into the operant chamber via a sliding rear wall within the delivery device, which also prevented either the experimental or social partner mouse from moving back into the mouse delivery device while the automatic door was open. Once the trial has ended, we removed the social partner mouse from the operant chamber through the side and placed the social partner mouse back into the 3D-printed mouse delivery device.

### Social defeat apparatus

The chronic social and witness defeat apparatus was adapted from previously published studies^26, 27^, schematically detailed in Fig. 3A. Defeats took place in transparent rectangular hamster cages (26.7 cm (w) × 48.3 cm (d) × 15.2 cm (h); Allentown, cat. no. PC10196HT) with paired steel-wire tops containing woodchip bedding. The hamster cage was divided in half by a clear perforated Plexiglas divider (0.6 cm (w) × 45.7 cm (d) × 15.2 cm (h); Nationwide Plastics, custom order), which creates a physical separation allowing for both physical (male) and witness (female) defeat procedures. The floor of the Allentown hamster cages was covered with clean woodchip bedding that we enriched with used home cage woodchip bedding to enhance territorial behavior.

### General operant self-administration experimental procedures

#### Social self-administration

The social self-administration procedure is based on previous studies^16, 17^, with some modifications. First, prior to any training sessions for operant access to a same-sex conspecific social partner, we gave the experimental C57Bl/6J mice three 5-min “magazine” training sessions for access to their partner, separated by 15-min each, in their operant chamber. Each session began with the presentation of the social partner-paired house light followed 10-s later by both a 2-s tone cue and the immediate insertion of an C57Bl/6J partner mouse; the house light remained on for the duration of the session to serve as a social partner-paired discriminative stimulus for the experimental C57Bl/6J mouse.

Next, we trained the experimental C57Bl/6J mice to self-administer access to their social same-sex partners during ten 48-min daily sessions using a discrete-trial design. Each 48-min session included twelve 4-min trials, schematically detailed in Figure 2A. The onset of the trials was signaled by the illumination of the social partner-paired house light, followed 10-s later by the insertion of the social partner-paired active lever; we allowed the experimental C57Bl/6J mice a maximum of 60-s to press the active lever on an FR1 reinforcement schedule before the lever automatically retracted.

Successful lever presses resulted in retraction of the active lever, followed first by a discrete 2-s tone cue and then the opening of the automatic guillotine door, through which their same-sex social partner was presented. The social partner-paired house light remained illuminated for 120-s, such that it terminated 110-s after the insertion of the active lever. We allowed the experimental C57Bl/6J mice access to their social partners until the house light turned off, at which point we removed their social partner through the main chamber door and returned the partner to the delivery device. After the termination of the social partner-paired house light, a 120-s inter-trial interval elapsed before the start of the next trial. We recorded the number of successful trials, the latency for active-lever press, and the number of inactive-lever presses. We trained two independent observers to identify non-affiliative social behaviors, like attack behavior, using previously operationalized metrics^17, 67^. If observed, these experimental mice were removed from the cohort.

#### Tests for social reward seeking under extinction conditions

We tested all experimental mice for social reward seeking (operationally defined as active-lever responding under non-reinforced extinction conditions) in 60-min test sessions. The social partner-paired house light (discriminative stimulus) signaled the start of the session. The social partner-paired active lever was inserted 10-s later. Active-lever presses caused the onset of the social partner-paired conditioned cue, with a 20-s fixed-interval period between cue presentations, but no social partner delivery. At the end of the 60-min session, the active lever retracted, and the house light was turned off. We recorded the number of social partner-paired cue presentations, as well as total active-lever and inactive-lever presses made.

#### Test for progressive-ratio reinforcement

We conducted all progressive-ratio tests using the same parameters we used for self-administration training, except for the trial design. Specifically, each progressive-ratio session began with the illumination of the social partner-paired house light, and 10-s later, the extension of the social partner-paired active lever. When the mouse met the number of active-lever presses required for a reinforced response, the active lever retracted and the 2-s cue was presented, followed by social reward delivery. The automatic door was opened for 10-s and the social partner was presented. Immediately after the automatic door closed, the active lever was re-extended. We removed the social partner mice immediately after 60-s of access. The progressive-ratio session was terminated if no reinforced responses occurred for 30-min or if a total duration of 2-h had elapsed. During the sessions, we increased the ratio of responses per social reward per the following sequence: 2, 4, 6, 9, 12, 15, 20, 25, 32, 40, 50, 62, 77, 95, 118, 145, 178, 219, 268, 328, 402, 492, 603, etc.^68^. The final completed response ratio represents the breakpoint value. <u>General social defeat and witness stress experimental procedures</u>

#### CD-1 screening using resident-intruder confrontations

We determined baseline unconditioned aggression in CD-1 mice by analyzing attack behavior in a variant of the resident-intruder task^24, 26, 69^. We screened CD-1 mice for aggression, and therefore inclusion as residents in the subsequent social defeat stress experiments, by performing 10-min once-daily resident-intruder tests for 3 consecutive days under dim -light conditions; the intruder mouse was always a novel unfamiliar C57Bl/6J mouse. Individual CD-1 mice were placed in an Allentown hamster cage that contained soiled bedding from their home cage. Aggression was scored on a scale from 0-3 (0 = no aggression; 1 = mild aggression defined by brief mounting and/or grooming; 2 = moderate aggression defined by several bouts of mounting and aggressive grooming with infrequent attack bouts; 3 = high aggression defined by frequent attack bouts). CD-1 mice with an average score of 2-3 were included as residents in subsequent social defeat experiments.

#### General chronic social and witness defeat stress procedures

The chronic social defeat and witness stress procedure is based on previous studies^26, 27^, with some modifications. First, chronic social defeat (male mice) and witness defeat (female mice) were run concurrently in the same experimental procedure, such that experimental female mice witnessed the physical defeat of experimental male mice. Prior to each daily defeat session, experimental male and female mice were acclimated to the defeat apparatus for 30-min. Experimental C57BL/6J mice were then subjected to a novel CD-1 aggressor mouse for 10 minutes once per day, over 10 consecutive days, on their designated side of the perforated divider. Experimental female C57Bl/6J mice observed these defeats from the opposite side of the perforated divider, allowing full sensory, but no physical, contact. Following the 10 minutes of agonistic interaction, the experimental mice were removed from the hamster cage to their pair-housed home cages. Control mice were pair housed throughout the defeat duration but never exposed to aggressive CD-1 mice. All mice were pair-housed for at least 4 hours between defeat sessions and operant self-administration sessions.

#### Hierarchical Clustering Analysis

In order to identify putative stress ‘susceptible’ and ‘resilient’ phenotypes of operant social behavior among mice, we used a standard hierarchical clustering algorithm (Ward’s method) toclassify mice into two clusters based on z-scored operant metrics from OSS. Five OSS metrics were chosen as features for analysis: 1) the difference in mean social rewards obtained over the last 3 days of training (Phase I) and stress exposure (Phase II), 2) active lever presses from the pre-stress social reward seeking test, 3) mean social rewards obtained from the last 3 days of self-administration testing during stress exposure, 4) active lever presses from the post-stress social reward seeking test, and 5) mean breakpoint from progressive ratio tests. These metrics were z-scored across experimental groups separately for male and female mice, and were selected based on their predicted relevance to each animal’s expression of socially motivated behavior following stress. The clustering results were scored and compared using the Calinski-Harabasz criterion, and distributions of OSS features for mice assigned to each cluster were tested for similarity using Kolmogorov-Smirnov tests. For population density comparisons, frequency distributions of OSS features were fitted with Gaussian Mixture Models and compared by calculating the overlap coefficient between them. See “General statistical analyses” below for more details. Clustering analyses and overlap coefficient calculations were performed using custom Python scripts.

#### Social index score

The five features used for ‘Hierarchical Clustering Analysis’ were aggregated to compose a summarizing measurement of operant social behavior for each mouse as their ‘social index’ score. We first compared the social index distributions for males and females based on their experimental condition (’control’, ‘stress’) to examine their similarity (overlap coefficient and Kolmogorov-Smirnov test; see ‘General statistical analyses” for details). We then plotted and examined the distributions of the raw OSS metrics for correlations with social index score using linear regressions, to study their relationship with the social index distribution throughout OSS. To determine which features were most impactful for males and females during the procedure, we used PCA (3-component) on the z-scored features that composed the social index and visually labeled each animal by their place on a color bar normalized to the social index range (males: −5 to 10; females: −5 to 5). The coefficients for each PC were reported (Pearson’s *r*) to show their correlations to each feature.

In order to examine the stability of social index throughout OSS, we compared the overall social index scores with those derived from using just 4 (leaving out PR) and 3 (leaving out PR and post-stress reward seeking) features. We used multiple regression analysis to determine how well feature spaces predicted the actual calculated social index scores (*r^2^*), and compared the best fit lines to the model that included all features using ANCOVA comparing their slope and intercept. We then looked at the relative changes in social index rank ordering between number of features used. Ranks were obtained by ordering scores from smallest to largest value for males and females. The similarity between ranks was examined using Kendall’s tau test. Feature scaling and PCA were performed using custom Python scripts, other statistical comparisons were performed as described in ‘General statistical analyses’.

#### General statistical analyses

For analysis of operant behavior, 2-way and 3-way ANOVAs were performed in Prism 8 (GraphPad). Significant main effects and interaction effects (p<0.05, two-tailed) were followed up with post-hoc tests (Fisher PLSD). Because our multifactorial ANOVAs yielded multiple main and interaction effects, we only report significant effects that are critical for data interpretation, unless specified otherwise in the text. Significant results of post-hoc analyses are indicated by asterisks in the figures. Continuous distributions were generated by fitting a Gaussian Mixture Model to the underlying frequency distribution, using bin sizes of 1 and 0.5 for social index and individual feature comparisons, respectively. Underlying discrete distributions and rank orders were statistically compared using Kolmogorov-Smirnov and Kendall’s tau tests in Python, respectively. Unless noted otherwise, overlap coefficients and curve-fitting was done using custom Python scripts. All other statistical tests were performed in GraphPad Prism 8. In Table S1 we provide a complete report of the relevant statistical results.

---

**Table S1.** Summary of statistical results.

| Figure number | Factor name | Test | F-value | p-value |
| --- | --- | --- | --- | --- |
| Figure 2B. Rewarded trials | Acquisition, between | 3way | $F_{1,88}=76.8$ | <0.0001**** |
| Figure 2B. Rewarded trials | Sex*Acquisition | 3way | $F_{1,88}=0.475$ | 0.4925 |
| Figure 2B. Rewarded trials (D8-10) mean | Acquisition, between | 2way | $F_{1,88}=130.7$ | <0.0001**** |
| Figure 2B. Rewarded trials (D8-10) mean | Acquisition * S-ex, between | 2way | $F_{1,88}=0.112$ | 0.7393 |
| Figure 2C. Lever presses | Acquisition * Lever | 3way | $F_{1,176}=14.33$ | 0.0002*** |
| Figure 2C. Lever presses | Acquisition * Lever * Sex | 3way | $F_{1,176}=1.66$ | 0.1985 |
| Figure 2C. Active lever presses | Acquisition, between, active lever males | 3way | $F_{1,176}=14.33$ | 0.001** |
| Figure 2C. Active lever presses | Acquisition, between, active lever females | 3way | $F_{1,176}=1.66$ | 0.0002*** |
| Figure 2D. Rewarded trials | Acquisition * Coat * Day | 3way | $F_{9,360}=1.61$ | 0.1115 |
| Figure 2D. Rewarded trials (D8-10) mean | Coat, between, acquirers | 3way | $F_{1,40}=2.71$ | 0.0226* |
| Figure 2D. Lever presses | Acquisition * Color * Lever, between | 3way | $F_{1,80}=0.855$ | 0.41 |
| Figure 2D. Active lever presses | Acquisition, between, 'same' group | 3way | $F_{1,80}=33.9$ | 0.0001*** |
| Figure 2E. Rewards obtained | Familiarity * Acquisition | 3way | $F_{1,39}=0.94$ | 0.339 |
| Figure 2E. Rewards obtained | Familiarity * Acquisition * Day | 3way | $F_{4,156}=0.22$ | 0.929 |
| Figure 3B. Rewarded trials | Condition, between, males | 2way | $F_{1,42}=6.08$ | 0.018* |
| Figure 3B. Rewarded trials (D8-10) mean | Condition, between, males | 2way | $F_{1,42}=26.7$ | <0.0001**** |
| Figure 3B. Lever presses | Condition * Lever, between, males | 2way | $F_{1,84}=12.1$ | 0.0008*** |
| Figure 3B. Active lever presses | Condition, between, males stress v control | 2way | $F_{1,84}=12.1$ | <0.0001**** |
| Figure 3B. PR | Condition, between, males | 2way | $F_{1,84}=8.46$ | 0.0058** |
| Figure 3C. Rewarded trials | Condition, between, females | 2way | $F_{1,16}=0.004$ | 0.9484 |
| Figure 3C. Rewarded trials (D8-10) mean | Condition, between, females | 2way | $F_{1,16}=0.0899$ | 0.7682 |
| Figure 3C. Active lever presses | Condition * Lever, between, females | 2way | $F_{1,32}=6.43$ | 0.018* |
| Figure 3C. Lever presses | Condition, between, females stress v control | 2way | $F_{1,32}=6.43$ | 0.0062** |
| Figure 3C. PR | Condition, between, females | 2way | $F_{1,17}=4.46$ | 0.0499* |
| Figure 4D. Distribution of z-score, males | Feature 'a' | OVLap<br>K-S | 57%<br>0.56 | 0.00133** |
| Figure 4D. Distribution of z-score, males | Feature 'b' | OVLap<br>K-S | 46%<br>0.68 | <0.0001**** |
| Figure 4D. Distribution of z-score, males | Feature 'c' | OVLap<br>K-S | 14%<br>0.91 | <0.0001**** |
| Figure 4D. Distribution of z-score, males | Feature 'd' | OVLap<br>K-S | 33%<br>0.75 | <0.0001**** |
| Figure 4D. Distribution of z-score, males | Feature 'e' | OVLap<br>K-S | 38%<br>0.66 | <0.0001**** |
| Figure 4D. Distribution of z-score, females | Feature 'a' | OVLap<br>K-S | 60%<br>0.56 | 0.126 |
| Figure 4D. Distribution of z-score, females | Feature 'b' | OVLap<br>K-S | 58%<br>0.56 | 0.126 |
| Figure 4D. Distribution of z-score, females | Feature 'c' | OVLap<br>K-S | 36%<br>0.56 | 0.126 |
| Figure 4D. Distribution of z-score, females | Feature 'd' | OVLap K-S | 21%<br>0.89 | 0.0007*** |
| Figure 4D. Distribution of z-score, females | Feature 'e' | OVLap K-S | 48%<br>0.56 | 0.126 |
| Figure 5B. Social index distribution, males | Control vs stress | OVLap K-S | 32%<br>0.74 | <0.0001**** |
| Figure 5B. Social index distribution, females | Control vs stress | OVLap K-S | 68%<br>0.56 | 0.126 |
| Figure 5C. Linear regression males | Social index * pre-stress self-administration | $r^2$ | 0.15 | 0.345 |
| Figure 5C. Linear regression males | Social index * pre-stress reward seeking | $r^2$ | 0.59 | <0.0001**** |
| Figure 5C. Linear regression males | Social index * post-stress self-administration | $r^2$ | 0.92 | <0.0001**** |
| Figure 5C. Linear regression males | Social index * post-stress reward seeking | $r^2$ | 0.82 | <0.0001**** |
| Figure 5C. Linear regression males | Social index * PR | $r^2$ | 0.85 | <0.0001**** |
| Figure 5D. PC1 correlation, males | Feature 'a' | r | 0.35 |  |
| Figure 5D. PC1 correlation, males | Feature 'b' | r | 0.33 |  |
| Figure 5D. PC1 correlation, males | Feature 'c' | r | 0.41 |  |
| Figure 5D. PC1 correlation, males | Feature 'd' | r | 0.6 |  |
| Figure 5D. PC1 correlation, males | Feature 'e' | r | 0.48 |  |
| Figure 5D. PC2 correlation, males | Feature 'a' | r | -0.5 |  |
| Figure 5D. PC2 correlation, males | Feature 'b' | r | 0.85 |  |
| Figure 5D. PC2 correlation, males | Feature 'c' | r | -0.07 |  |
| Figure 5D. PC2 correlation, males | Feature 'd' | r | -0.16 |  |
| Figure 5D. PC2 correlation, males | Feature 'e' | r | 0.05 |  |
| Figure 5E. Linear regression females | Social index * pre-stress self-administration | $r^2$ | 0.43 | 0.08 |
| Figure 5E. Linear regression females | Social index * pre-stress reward seeking | $r^2$ | 0.72 | 0.0008*** |
| Figure 5E. Linear regression females | Social index * post-stress self-administration | $r^2$ | 0.87 | <0.0001**** |
| Figure 5E. Linear regression females | Social index * post-stress reward seeking | $r^2$ | 0.78 | 0.00013*** |
| Figure 5E. Linear regression females | Social index * PR | $r^2$ | 0.54 | 0.0199* |
| Figure 5F. PC1 correlation, females | Feature 'a' | r | 0.23 |  |
| Figure 5F. PC1 correlation, females | Feature 'b' | r | 0.36 |  |
| Figure 5F. PC1 correlation, females | Feature 'c' | r | 0.51 |  |
| Figure 5F. PC1 correlation, females | Feature 'd' | r | 0.69 |  |
| Figure 5F. PC1 correlation, females | Feature 'e' | r | 0.29 |  |
| Figure 5F. PC2 correlation, females | Feature 'a' | r | 0.78 |  |
| Figure 5F. PC2 correlation, females | Feature 'b' | r | -0.33 |  |
| Figure 5F. PC2 correlation, females | Feature 'c' | r | 0.21 |  |
| Figure 5F. PC2 correlation, females | Feature 'd' | r | -0.04 |  |
| Figure 5F. PC2 correlation, females | Feature 'e' | r | -0.05 |  |
| Figure 6B. Regression 5 features males | Actual vs predicted social index scores | $r^2$ | 1 | <0.0001**** |
| Figure 6B. Regression 4 features males | Actual vs predicted social index scores | $r^2$ | 0.98 | <0.0001**** |
| Figure 6B. Regression 3 features males | Actual vs predicted social index scores | $r^2$ | 0.89 | <0.0001**** |
| Figure 6B. Regressions comparison, males | Slopes | ANCOVA | $F_{2, 10} = 2.801$ | 0.065 |
| | Intercepts | ANCOVA | $F_{2, 110} = 1.8e^{-12}$ | 0.9999 |
| Figure 6C. Regression 5 features females | Actual vs predicted social index scores | $r^2$ | 1 | <0.0001**** |
| Figure 6C. Regression 4 features females | Actual vs predicted social index scores | $r^2$ | 0.94 | <0.0001**** |
| Figure 6C. Regression 3 features females | Actual vs predicted social index scores | $r^2$ | 0.88 | <0.0001**** |
| Figure 6C. Regressions comparison, females | Slopes | ANCOVA | $F_{2, 48}=2.801$ | 0.361 |
| | Intercepts | ANCOVA | $F_{2, 50}=3.5e^{-13}$ | 0.9999 |
| Figure 6D. Rank similarity, males | 5 v 4 features | Kendall's tau | 0.9 | <0.0001**** |
|  | 5 v 3 features | Kendall's tau | 0.82 | <0.0001**** |
| Figure 6D. Rank similarity, females | 5 v 4 features | Kendall's tau | 0.91 | <0.0001**** |
|  | 5 v 3 features | Kendall's tau | 0.8 | <0.0001**** |
| Figure S1B. Rewarded trials | Acquisition, between | 3way | $F_{1,36}=37.6$ | <0.0001**** |
| Figure S1B. Rewarded trials (D8-10) mean | *Acquisition \* Experience, between* | 2way | $F_{1,36}=3.01$ | 0.0912 |
| Figure S1B. Rewarded trials (D8-10) mean | Acquisition * Experience, between acquirers | 2way | $F_{1,36}=3.01$ | 0.0003*** |
| Figure S1C. Lever presses | *Acquisition \* Experience \* Lever* | 3way | $F_{1,72}=1.12$ | 0.294 |
| Figure S1C. Lever presses | Acquisition * Lever, between naïve | 3way | $F_{1,72}=25.47$ | 0.0008*** |
| Figure S1C. Lever presses | Acquisition * Lever, between experienced | 3way | $F_{1,72}=25.47$ | <0.0001**** |
| Figure S1C. Active lever presses | Acquisition * Experience, between acquirers | 3way | $F_{1,72}=1.19$ | 0.036* |

